# Organization and replicon interactions within the highly segmented genome of *Borrelia burgdorferi*

**DOI:** 10.1101/2023.03.19.532819

**Authors:** Zhongqing Ren, Constantin N. Takacs, Hugo B. Brandão, Christine Jacobs-Wagner, Xindan Wang

## Abstract

*Borrelia burgdorferi*, a causative agent of Lyme disease, contains the most segmented bacterial genome known to date, with one linear chromosome and over twenty plasmids. How this unusually complex genome is organized, and whether and how the different replicons interact are unclear. We recently demonstrated that *B. burgdorferi* is polyploid and that the copies of the chromosome and plasmids are regularly spaced in each cell, which is critical for faithful segregation of the genome to daughter cells. Regular spacing of the chromosome is controlled by two separate partitioning systems that involve the protein pairs ParA/ParZ and ParB/SMC. Here, using chromosome conformation capture (Hi-C), we characterized the organization of the *B. burgdorferi* genome and the interactions between the replicons. We uncovered that although the linear chromosome lacks contacts between the two replication arms, the two telomeres are in frequent contact. Moreover, several plasmids specifically interact with the chromosome *oriC* region, and a subset of plasmids interact with each other more than with others. We found that SMC and the SMC-like MksB protein mediate long-range interactions on the chromosome, but they minimally affect plasmid-chromosome or plasmid-plasmid interactions. Finally, we found that disruption of the two partition systems leads to chromosome restructuring, correlating with the mis-positioning of chromosome *oriC*. Altogether, this study revealed the conformation of a complex genome and analyzed the contribution of the partition systems and SMC family proteins to this organization. This work expands the understanding of the organization and maintenance of multipartite bacterial genomes.

## Author summary

Genomes are highly organized in cells to facilitate biological processes. *Borrelia burgdorferi*, an agent of Lyme disease, carries one linear chromosome and more than twenty plasmids, in what is known as one of the most segmented bacterial genomes. How the different replicons interact with each other is unclear. Here we investigate the organization of this highly segmented genome and the protein factors that contribute to this organization. Using chromosome conformation capture assays, we determined the interactions within the chromosome, between chromosome and plasmids, and between the plasmids. We found that the two telomeres of the chromosome interact with each other; a subset of plasmids interact with the chromosomal replication origin region; and a subset of plasmids preferentially interact with one another. Finally, we revealed that two structural maintenance of chromosomes family proteins, SMC and MksB, promote long-range DNA interactions on the chromosome, and the two partition systems, ParA/ParZ and ParB/SMC, contribute to chromosome structure. Altogether, we characterized the conformation of a highly segmented genome and investigated the functions of different genome organizers. Our study advances the understanding of the organization of highly segmented bacterial genomes.

## Introduction

*Borrelia burgdorferi* causes Lyme disease, the most prevalent vector-borne infectious disease in Europe and North America [1, 2]. Although the *B. burgdorferi* genome is only ∼1.5 megabasepairs in size, it includes one linear chromosome and more than 20 plasmids (circular and linear) and is, to our knowledge, the most segmented bacterial genome [3–6]. Recently, using fluorescence microscopy to visualize loci on the chromosome and 16 plasmids, we found that *B. burgdorferi* contains multiple copies of its genome segments *per* cell, with each copy regularly spaced along the cell length [7].

In bacteria, the broadly conserved *parABS* partitioning system plays an important role in the segregation of chromosome and plasmids [8–15]. ParA dimerizes upon ATP binding and non-specifically binds to the DNA [16–19]. Centromeric ParB proteins bind to the *parS* sequences scattered around the origin of replication and spread several kilobases to nearby regions, forming a nucleoprotein complex [20–25]. The ParB-DNA nucleoprotein complex interacts with DNA-bound ParA-ATP dimers and stimulates the ATPase activity of ParA, leading to the release of ParA from the DNA and the formation of a ParA concentration gradient along the nucleoid [12, 15, 17, 26]. It is thought that repeated cycles of ParA and ParB interaction and release, together with the translocating forces from elastic chromosome dynamics [27–30] or the chemical ParA gradient [31, 32], promote the segregation of the two newly replicated ParB-origin complexes from one another [27, 29]. In addition, ParB plays a separate role in recruiting the broadly conserved SMC complex onto the chromosomal origin region [13, 14]. Once loaded, SMC moves away from the loading sites and typically tethers the two replication arms together, facilitating the resolution and segregation of the two sister chromosomes [33–35].

We discovered that in *B. burgdorferi*, the segregation and positioning of the multicopy chromosomal origins of replication (*oriC*) require the concerted actions of the ParB/SMC system and a newly discovered ParA/ParZ system [7]. ParZ, a centromere-binding protein, substitutes ParB to work with ParA and plays a major role in chromosome segregation [7]. Although *B. burgdorferi* ParB does not appear to partner with ParA, it is still required to recruit SMC to *oriC*. SMC in turn contributes to *oriC* positioning [7]. Overall, these findings advanced our understanding of *oriC* segregation in *B. burgdorferi*. However, the information on the organization of the bulk of the chromosome and the interactions among the various genome segments in this bacterium is still lacking.

Chromosome conformation capture assays (Hi-C) have significantly advanced our understanding of bacterial genome folding and interactions [34, 36–41]. Along bacterial genomes, short-range self-interacting domains called chromosome interaction domains (CIDs) have been observed and are shown to be dictated mostly by transcription, with domain boundaries correlating with highly transcribed genes. In bacteria that contain the canonical SMC complex, the two replication arms of the chromosome are juxtaposed together, whereas bacteria that only encode SMC-like MukBEF and MksBEF analogs do not show inter-arm interactions [37, 39].

More recent efforts have begun to reveal the genome conformation of bacteria containing multiple replicons. In *Agrobacterium tumefaciens*, the origins of the four replicons are clustered together, which regulates DNA replication and drives the maintenance of this multipartite genome [41, 42]. Similarly, the two origins of *Brucella melitensis* chromosomes also showed frequent interactions [43]. In *Vibrio cholerae*, the origin of Chromosome 2 (Ch2) interacts with the *crtS* region on Chromosome 1 (Ch1) for replication control, and the terminus region of Ch1 and Ch2 are interacting for coordinated replication termination and terminus segregation [40, 44]. These findings suggest that multipartite genomes harness inter-replicon interactions as a mechanism for replication regulation and genome maintenance. In this study, we aimed at understanding how *B. burgdorferi* organizes its ∼20 replicons and how the partitioning proteins and SMC homologues contribute to genome organization.

## Results

### The organization of the linear *B. burgdorferi* chromosome

To determine the organization of the highly segmented genome of *B. burgdorferi*, we performed Hi-C on exponentially growing cultures of the infectious, transformable strain S9 (**Table S1** and **Fig. 1A, B**). After mapping the reads and plotting the data, we observed many white lines on the Hi-C map, especially in regions of the map corresponding to the plasmids (**Fig. 1B**). These white lines indicate the presence of repetitive sequences on the affected replicons, which were omitted during sequence mapping. The genome-wide Hi-C interaction map (**Fig. 1B**) has four distinct regions: an intra-chromosome interaction map in the lower left quadrant, a plasmid-chromosome interaction map with identical, mirrored copies in the top left and lower right quadrants, and a plasmid-plasmid interaction map in the top right quadrant. The chromosome displayed strong short-range interactions as evident by the primary diagonal (**Fig. 1B**, lower left quadrant).

**Figure 1.**
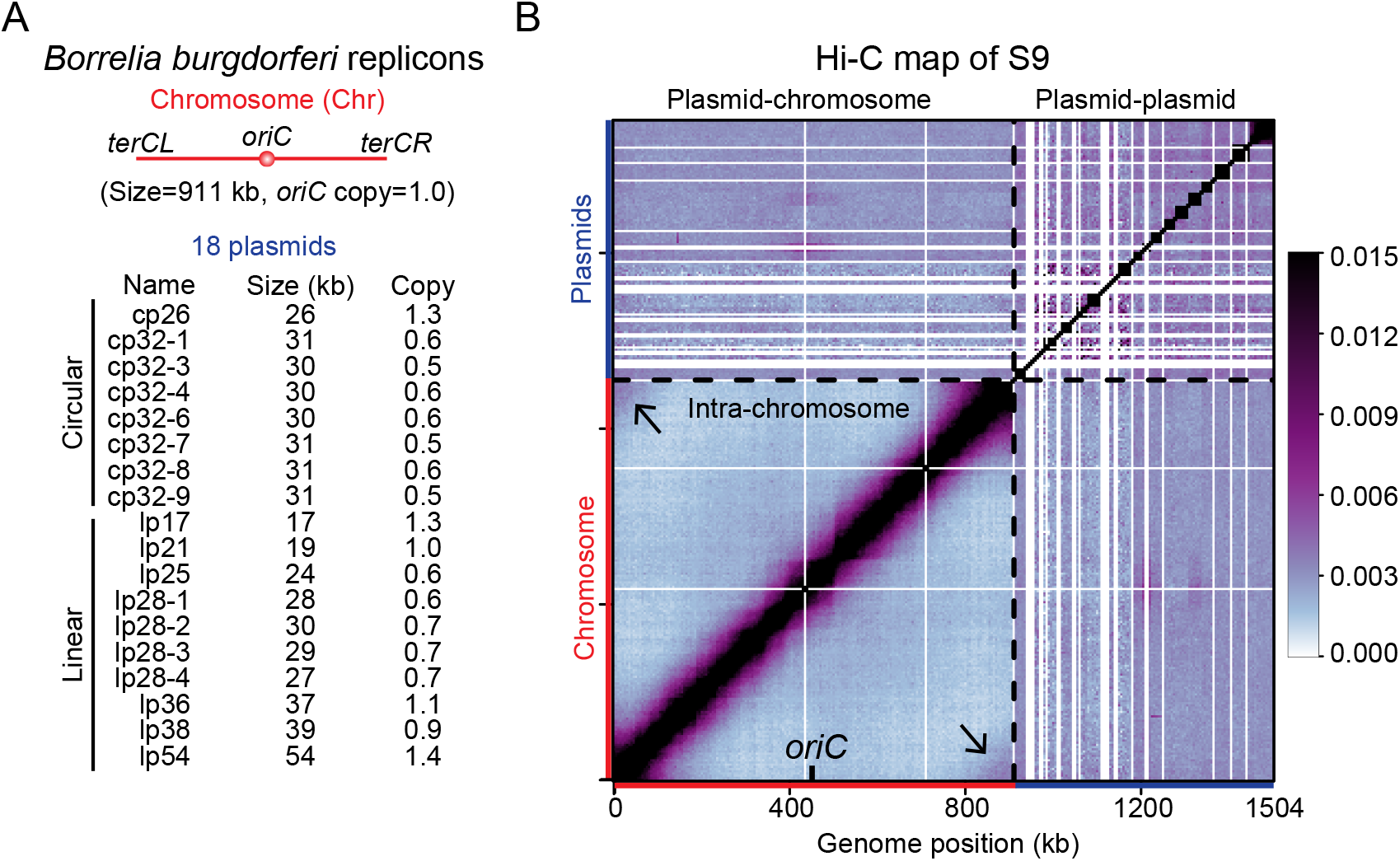
Genome-wide organization of *B. burgdorferi* replicons. **(A)** The *B. burgdorferi* S9 wild-type strain has a linear chromosome (Chr), 8 circular plasmids and 10 linear plasmids. The replication origin of the chromosome is labeled as *oriC*. The sizes (in kb) and relative copy numbers of the plasmids are listed. The relative copy number of each plasmid were previously measured using whole genome sequencing analysis [7], and is shown relative to the copy number of *oriC*. **(B)** Normalized Hi-C matrix showing interaction frequencies for pairs of 5-kb bins across the genome of *B. burgdorferi* S9. x and y-axes show genome positions. The chromosome and the plasmids are indicated by red and blue bars, respectively. *oriC* is labeled on the x-axis. The boundary between the chromosome and the plasmids are indicated by black dotted lines. The plasmids are ordered alphabetically from cp26 to lp54, from left to right and bottom to top, respectively. The whole map was divided into four regions: the bottom left region shows intra-chromosomal interactions, the top left and bottom right regions show plasmid-chromosome interactions, and the top right region represents plasmid-plasmid interactions. We used the same convention for all whole-genome Hi-C and Hi-C derivative plots in this study. The color scale depicting Hi-C interaction scores in arbitrary unit is shown at the right.

Interestingly, a secondary diagonal representing inter-arm interactions was absent from the Hi-C map. This was unexpected as *B. burgdorferi* encodes an SMC protein homolog and all SMC-carrying bacteria tested so far display chromosome with inter-arm interactions [34, 36, 38, 39, 41, 45, 46]. We note that although *B. burgdorferi* does contain a homolog of the ScpA subunit of the SMC complex, it does not encode the other subunit, ScpB [3]. Thus, the absence of the SMC-ScpAB holo-complex might explain the absence of chromosome arm alignment in *B. burgdorferi* (see Discussion). Additionally, the two ends of the chromosome, the left and right telomeres (*terCL* and *terCR*) displayed a striking interaction with each other (**Fig. 1B**, black arrows in lower left quadrant). Since *B. burgdorferi* is polyploid [7], it is unclear whether the interacting *terCL* and *terCR* are located on the same chromosome or on adjacent chromosome copies.

### Interactions between the chromosome and 18 plasmids

Qualitatively, plasmid-chromosome interactions were weaker than short-range interactions within the chromosome (i.e. the primary diagonal of the bottom left quadrant), but were stronger than long-range interactions within the chromosome (i.e. outside of the primary diagonal on the bottom left quadrant) (**Fig. 1B**). We plotted the distribution of these types of interaction frequencies and found that the differences were statistically significant (**Fig. 2**). To better show the plasmid-chromosome interactions, we analyzed the interaction of each plasmid with each 5-kb bin on the chromosome (**Fig. 3A**). Interestingly, a subset of the linear plasmids, namely lp17, lp21, lp25, and lp28-3, showed stronger interactions with the chromosomal origin region compared with the rest of the chromosome (**Fig. 3A**). These interactions are reminiscent of the origin clustering interactions mediated by centromeric proteins in *A. tumefaciens*, which are critical for the replication and maintenance of the secondary replicons in that bacterium [41, 42]. Notably, the plasmid-chromosome interactions observed here are weaker than those observed in *A. tumefaciens*, and only 4 out of 18 plasmids showed these specific interactions with the chromosome, thus the biological function of these interactions is unclear (see Discussion).

**Figure 2.**
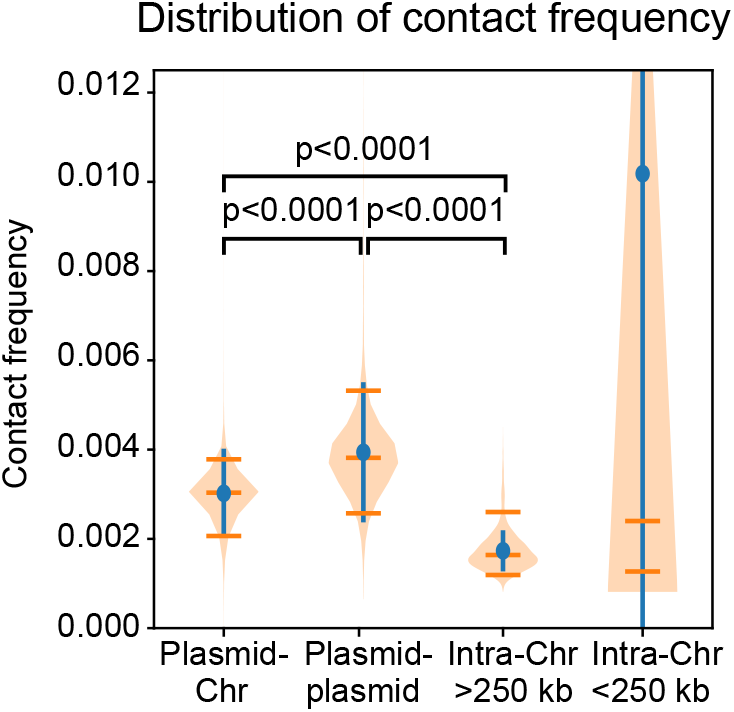
Hi-C contact frequencies for different types of interactions. Distributions of Hi-C contact frequencies measured for different types of interactions are shown as violin plots. Blue lines indicate standard deviations of the values. Orange lines indicate the median, 5^th^ and 95^th^ percentile of the data. The *p-*values were computed using a Mann-Whitney U test. All comparisons were done for data binned at 5 kb resolution.

**Figure 3.**
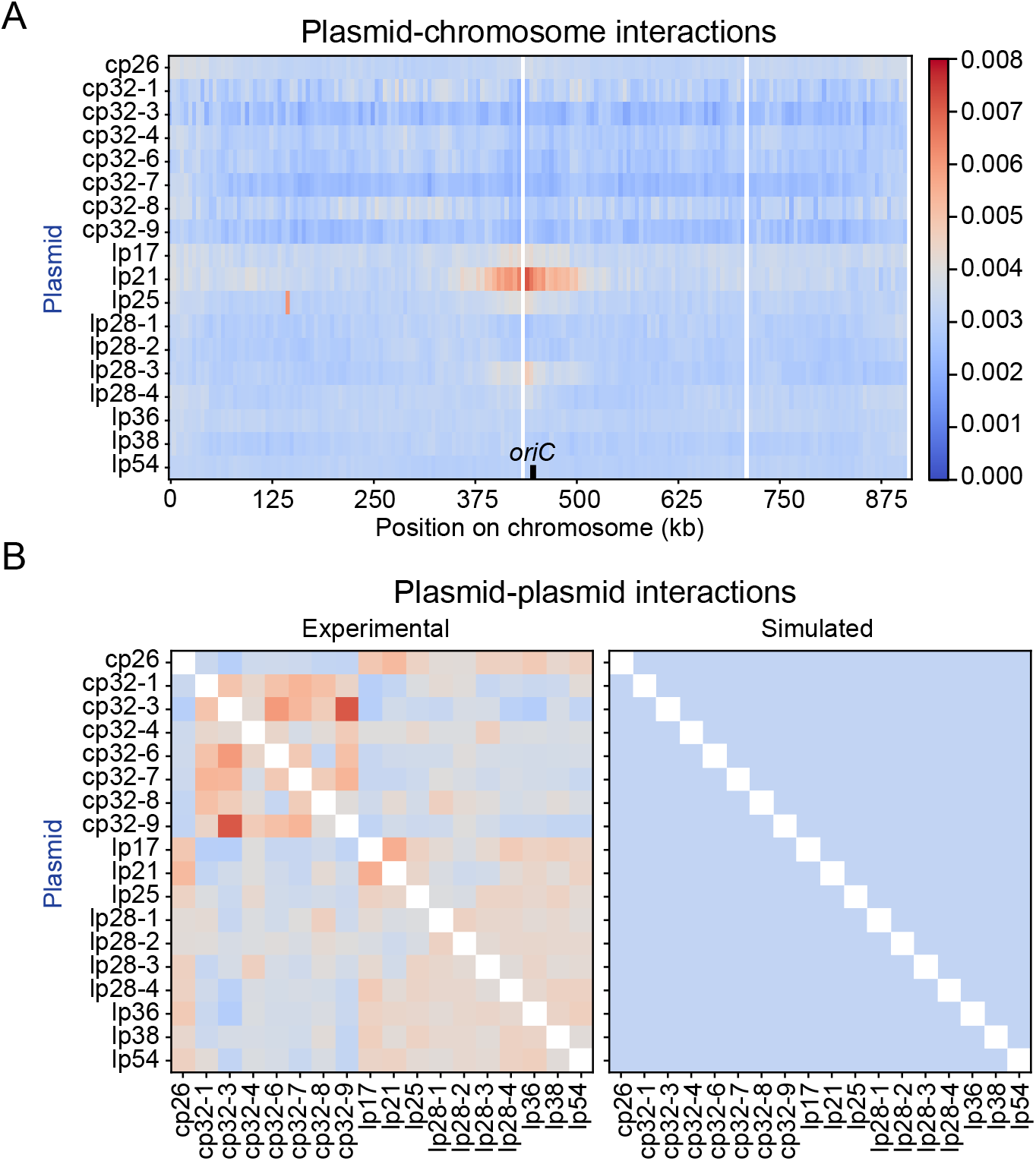
Plasmid-chromosome and plasmid-plasmid interactions. **(A)** The heatmap of plasmid interactions with chromosome loci in WT *B. burgdorferi* train S9. To generate the interaction score between each plasmid and each chromosome locus, the Hi-C interaction scores in consecutive bins are summed according to each plasmid. The plot shows averaged data of two replicates. The x-axis indicates the genome position on the chromosome. The y-axis specifies different plasmids. The color scale depicting interaction scores in arbitrary unit is shown at the right. The color scale depicting relative interaction frequency in arbitrary unit is shown at the right. **(B)** Left, the experimentally measured interaction frequencies between plasmids. To generate the interaction score within every pair of plasmids, the Hi-C interaction scores in consecutive bins are summed according to each plasmid. The data are normalized such that each row has the same total score. This normalization ignores the plasmid-chromosome interactions. The plot shows averaged data of two replicates. The x-axis and y-axis indicate the different plasmids of *B. burgdorferi* strain S9. the simulated interaction frequencies between plasmids based on plasmid copy number and plasmid sizes (see Materials and Methods). The normalization method is the same as the experimental data shown on the light. The color scale is the same as in **(A).** The simulated maps with iterative correction or in a finer color scale can be found in **Fig. S1**.

### Plasmid-plasmid interactions

Plasmid-plasmid interactions are depicted in the top right quadrant of the Hi-C map (**Fig. 1B**) and appeared stronger than plasmid-chromosome interactions (**Fig. 1B**, top left quadrant, and **Fig. 2**) and long-range interactions within the chromosome (**Fig. 1B**, outside of the primary diagonal on the bottom left quadrant, and **Fig. 2**). To better understand the interactions between every two plasmids, we recalculated the interaction frequencies after excluding the plasmid-chromosome interactions from the analysis (**Fig. 3B**). We note that the sizes of the 18 plasmids ranged from 17 kb to 54 kb [3, 4] and that their copy numbers had been previously determined by microscopy and whole genome sequencing, ranging from 0.5 to 1.3 relative to the copy number of the *oriC* locus [7] (**Fig. 1A**). To understand whether these sizes and copy numbers of the plasmids could impact plasmid-plasmid interactions, we used these numbers to simulate the plasmid-plasmid interaction frequencies, assuming that all the plasmids were freely diffusing in the cytoplasm (see Materials and Methods for simulation details). Our simulation showed that plasmids that have a bigger size or a higher copy number interacted more with other plasmids in the raw Hi-C maps before any corrections (**Fig. S1A, B, top panels**). However, these preferential interactions did not show up after our standard procedure of iterative corrections for the Hi-C maps [47] (**Fig. S1A, B**, middle panels), unless a very fine color scale was applied (**Fig. S1A, B**, bottom panels). Interestingly, in our experiment (**Fig. 3B**, left), the interactions among the seven cp32 plasmids (cp32-1, cp32-3, cp32-4, cp32-6, cp32-7, cp32-8, cp32-9) and among the other 11 plasmids were higher than expected for random encounters based on simulations (**Fig. 3B**, right). Thus, the preferential interactions between plasmids we observed in our experiment could not be explained solely by the size and copy number difference in the plasmids. Since repetitive sequences between different plasmids were removed during mapping, we believe that these higher-than-expected interactions observed in our experiment are genuine and not due to erroneous normalization or mapping. The molecular mechanism of plasmid-plasmid interactions remains to be determined.

### Clustering analysis of *smc* and *par* mutants

The highly conserved SMC family proteins and the DNA partitioning proteins are central players in bacterial chromosome organization and segregation [48, 49]. *B. burgdorferi* has a canonical SMC, encoded by gene *bb0045*, as well as an MksB protein, encoded by gene *bb0830*, but lacks the genes encoding the accessory proteins ScpB, MksE, and MksF [3]. Additionally, *B. burgdorferi* employs two partition systems for the positioning of its multicopy *oriC* loci, ParB/SMC and ParA/ParZ [7]. To understand the contribution of these factors to *B. burgdorferi* genome interactions, we performed Hi-C on a collection of mutants (**Table S1**). Essentially, the genes of interest were replaced with a gentamycin or kanamycin resistance gene. The control strain CJW_Bb284 had the gentamycin marker inserted in a non-coding region located in between the convergently-oriented *parZ* and *parB* genes, in the otherwise wild-type (WT) *parAZBS* locus. The Hi-C maps of strain CJW_Bb284 were almost identical to the maps generated using the parental WT strain S9 (**Fig. S2**). Additionally, our Hi-C experiments on WT, control, and every mutant were done in two biological replicates that showed nearly identical results (**Fig. S3**).

To compare the different mutants, we performed a clustering analysis using the contact probability curves of our 22 Hi-C samples so that mutants that had similar profiles of contact probabilities would be grouped together (**Fig. 4, S4**). Using the Silhouette method [50], we found that the mutants could be divided into six groups (**Fig. 4A, B**) (see Materials and Methods): group 1 includes WT and the control strain CJW_Bb284 (**Fig. 4B, C**, **Fig. S2**); group 2 includes *Δsmc* (**Fig. 4B, D**); group 3 includes *ΔmksB* (**Fig. 4B, E**); group 4 includes *ΔparB*, *ΔparS* and *ΔparBS* (**Fig. 4B, F)**; group 5 includes *ΔparA*, *ΔparZ* and *ΔparAZ* (**Fig. 4B, G**); and group 6 includes *ΔparAZBS* (**Fig. 4B, H**)

**Figure 4.**
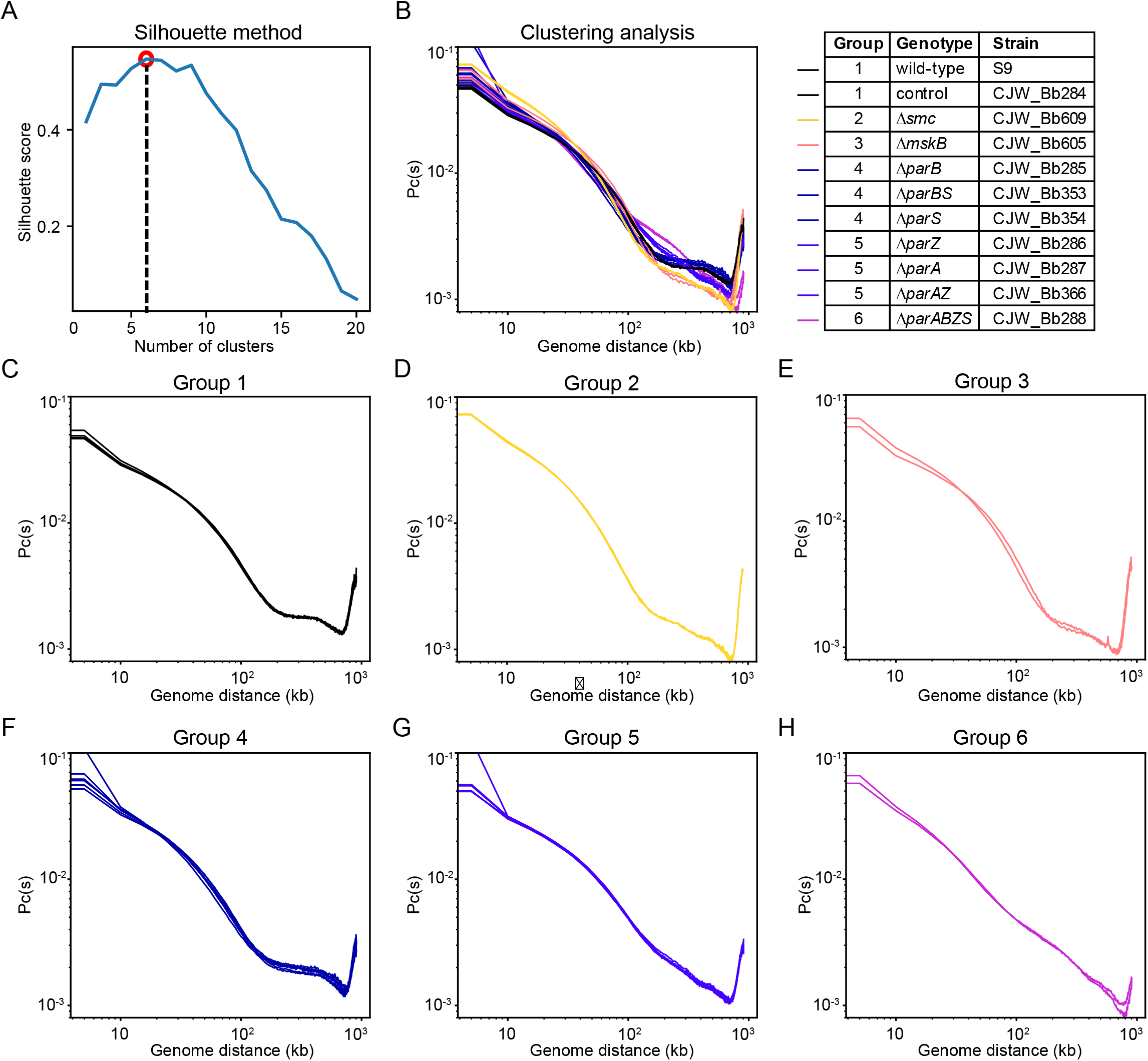
Clustering analysis of different mutants. **(A)** Determination of the optimal number of clusters of contact probability curves, Pc(s), for k-means clustering (see Materials and Methods). The number of clusters was determined by identifying the peak in Silhouette score. This analysis suggests six optimal groupings, which is indicated by the red circle and black dotted line. **(B)** Pc(s) curves of all the samples. Grouping results of the 11 strains are listed on the right. Two biological replicates of each strain are plotted. Individual Pc(s) curves can be found in **Fig. S4**. (**C-I**) Curves of the same group in (**B**) are plotted in different panels.

This grouping analysis based on Hi-C results indicates that the control strain CJW_Bb284 behaves the same as its parental WT strain; SMC and MksB have different effects on chromosome folding; ParB and *parS* work as a unit; ParA and ParZ work together; and ParB/*parS* and ParA/ParZ have additive effects because *ΔparAZBS* formed its own group. Notably, our recent ChIP-seq and microscopy analyses [7] have indicated that ParB binds to *parS* and recruits SMC to the origin region, and ParZ works with ParA; disrupting *parBS* barely changed *oriC* spacing; deleting *parA*, *parZ* or *parAZ* had similar effects and dramatically changed the even spacing of *oriC* in the polyploid cells; finally, deleting *parBS* and *parA* caused a stronger defect in *oriC* spacing than *ΔparAZ* alone [7]. Therefore, the grouping of mutants based on Hi-C analysis here (**Fig. 4B**) is largely consistent with our previous cytological characterization of these mutants [7]. This agreement reveals the robustness of our assays.

### SMC and MksB mediate long-range interactions within the chromosome

In our clustering analysis, the two biological replicates of *Δsmc* fell in one group (group 2) and replicates of *ΔmksB* fell into a separate group (group 3) (**Fig. 4B, D, E**). To understand how *Δsmc* and *ΔmksB* affect genome contacts, we analyzed the log_2_ ratios of the Hi-C maps between each mutant strain and the relevant control. (**Fig. 5A-F**). We observed that both *Δsmc* and *ΔmksB* strains had decreased long-range DNA contact compared with the control (**Fig. 5D-F**, blue pixels in black trapezoid). Specifically, as seen on the Hi-C contact probability decay curves (**Fig. 5G-I**), in *Δsmc*, loci separated by ∼60 kb or greater had decreased frequency of contacts compared with the control, and in *ΔmksB*, loci separated by ∼100 kb or greater had decreased frequency of contact compared with the control (**Fig. 5H, I**, black dotted lines). These data indicate that both SMC and MksB promote long-range DNA contacts and that their effects are different enough to fall into different groups in our clustering analysis. We noted that *B. burgdorferi* is missing the ScpB subunit of the SMC complex, as well as the MksE and MksF subunits of the MksBEF complex. However, previous work showed that purified *B. subtilis* SMC protein (in the absence of ScpA and ScpB) is able to form DNA loops *in vitro* [51]. Our results suggest that the incomplete SMC/Mks complexes may form DNA loops in *B. burgdorferi*. Curiously, the absence of SMC or MksB enhanced the *terCL-terCR* interactions (**Fig. 5E, F**, black arrows), suggesting that these proteins reduce the contacts between the telomeres. Finally, we note that both SMC and MksB mainly affect interactions within the chromosome and not between chromosome and plasmid or among the plasmids (**Fig. 5A-F, S5-7**).

**Figure 5.**
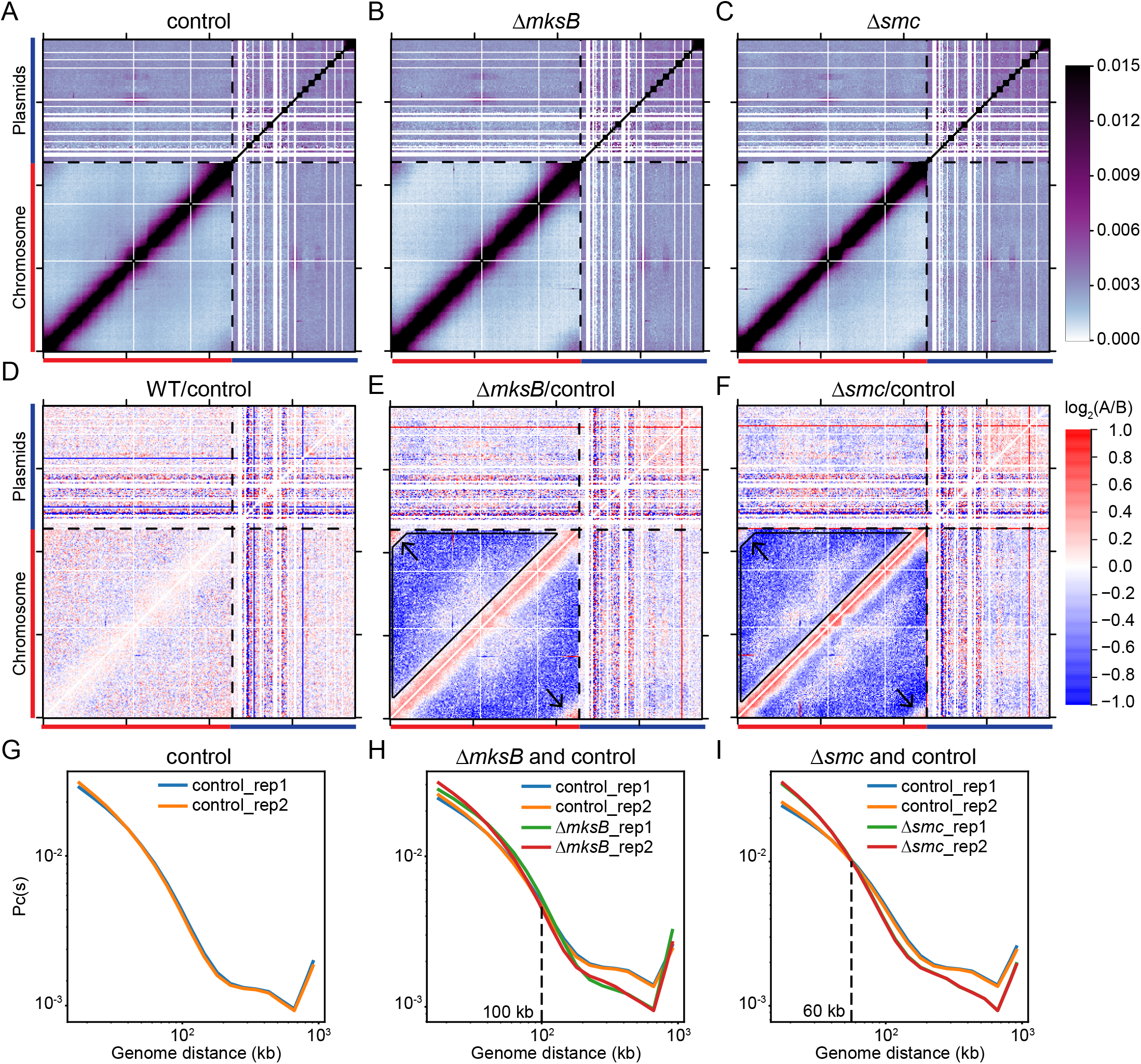
SMC and MksB mediate long-range DNA interactions. **(A-C)** Normalized Hi-C interaction maps of the control (CJW_Bb284), Δ*mksB* (CJW_Bb605,) and Δ*smc* (CJW_Bb609) strains. Black dotted lines mark the boundary between the depiction of the chromosome and that of the plasmids. The color scale depicting Hi-C interaction scores in arbitrary unit is shown at the right. **(D-F)** Log_2_ ratio plots comparing different Hi-C matrices. Log_2_(matrix 1/matrix 2) was calculated and plotted in the heatmaps. Matrix 1/ matrix 2 are shown at the top of each plot. The color scale is shown at the right of panel **(F)**. Black arrows point to *terCL-terCR* interactions. Black trapezoids indicate reduced interactions in the mutants. **(G-I)** Contact probability decay Pc(s) curves of indicated Hi-C matrices. Pc(s) curves show the average contact frequency between all pairs of loci on the chromosome separated by set distance (*s*). The x-axis indicates the genomic distance of separation in kb. The y-axis represents averaged contact frequency. The curves were computed for data binned at 5 kb. The intersection points of mutant and control curves are indicated by black dotted lines.

### Contribution of ParB/*parS* and ParA/ParZ to chromosome organization

In the grouping analysis, Δ*parS*, Δ*parB* and Δ*parBS* fell in the same group (group 4) (**Fig. 4B, F**), consistent with previous finding that ParB and *parS* act as a unit [7]. The absence of *parB* and/or *parS* caused similar changes to genome interactions compared with the control (**Fig. 6A-F**): *terCL-terCR* interactions decreased (**Fig. 6D-F**, blue pixels indicated by black arrows); longer range (>150 kb) interactions within the chromosome increased (**Fig. 6D-F**, red pixels within black trapezoid); and short-range interactions (50-150 kb) decreased (**Fig. 6D-F**, blue pixels between black trapezoid and the red line). These trends are opposite to those observed in *Δsmc* or *ΔmksB* (**Fig. 5E, F**). Since ParB recruits SMC to the *oriC* region in *B. burgdorferi* [7], the loss of *parBS* could lead to increased non-specific loading of SMC on the chromosome. Thus, these results are consistent with a scenario in which non-specific loading of SMC to the chromosome outside of the *oriC* region (i.e. independent of ParB/*parS*) is the major contributor to long-range chromosome interactions.

**Figure 6.**
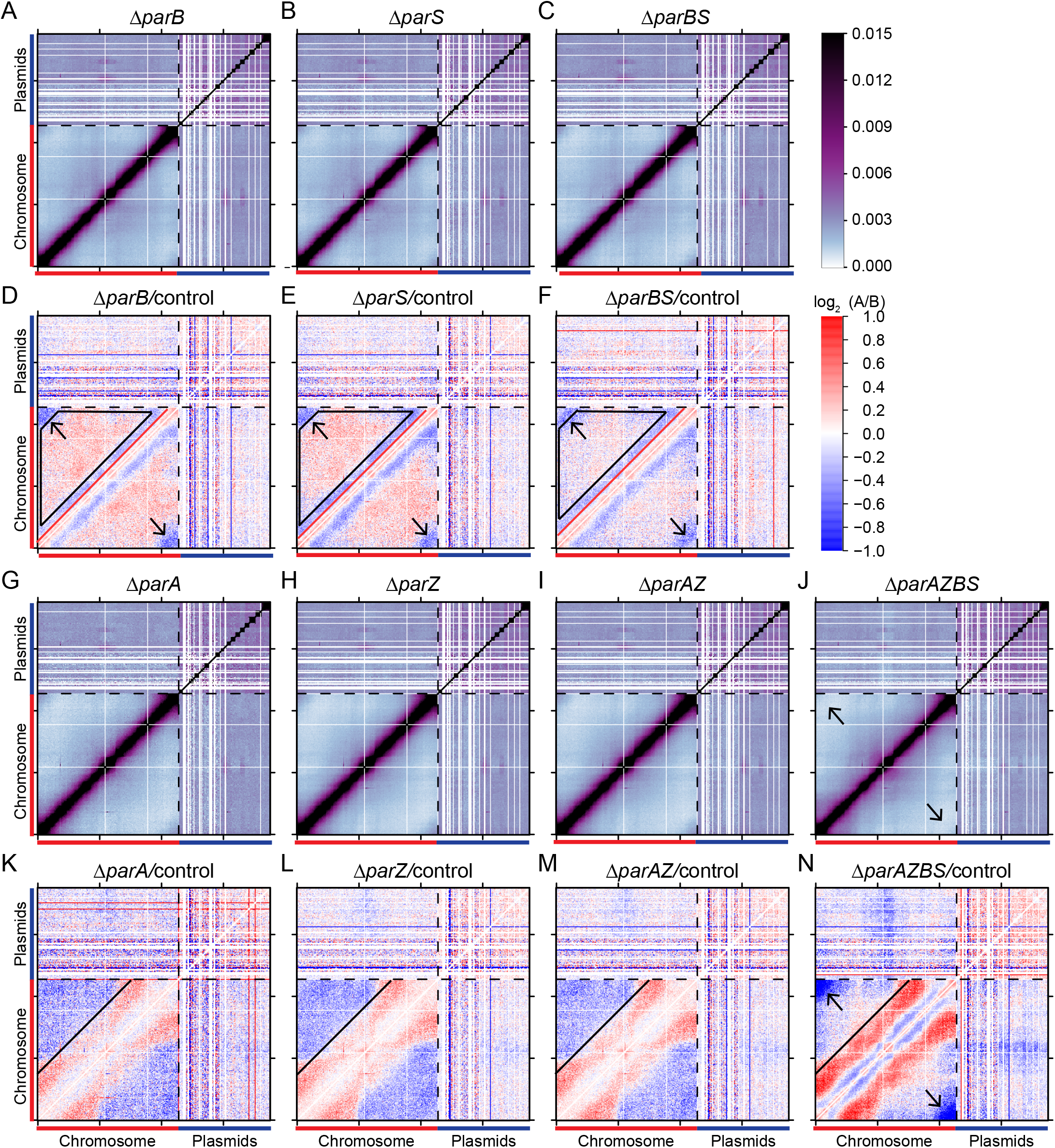
Disruption of the partition systems re-structures the genome. **(A-C)** Normalized Hi-C interaction maps of the Δ*parB* (CJW_Bb353), Δ*parS* (CJW_Bb354), and Δ*parBS* (CJW_Bb285) strains. Black dotted lines indicate the boundary between the chromosome and the plasmids. The color scale depicting Hi-C interaction scores in arbitrary unit is shown at the right. **(D-F)** Log_2_ ratio plots comparing Δ*parB* (CJW_Bb353), Δ*parS* (CJW_Bb354), and Δ*parBS* (CJW_Bb285), respectively, with the control (CJW_Bb284) strain. Black arrows point to blue pixels *terCL-terCR* interactions. Black trapezoids indicate area of read pixels. Red lines indicate the boundary between red and blue pixels. The color scale is shown at the right. **(G-J)** Normalized Hi-C interaction maps of the Δ*parA* (CJW_Bb366), Δ*parZ* (CJW_Bb286), Δ*parAZ* (CJW_Bb287) and Δ*parAZBS* (CJW_Bb288) strains. Black arrows indicate *terCL-terCR* interactions. **(I-N)** Log_2_ ratio plots comparing Δ*parA* (CJW_Bb366), Δ*parZ* (CJW_Bb286), Δ*parAZ* (CJW_Bb287), or Δ*parAZBS* (CJW_Bb288) with the control (CJW_Bb284) strain. Solid black lines indicate the boundary between red and blue pixels. Black arrows indicate *terCL-terCR* interactions.

Group 5 contains *ΔparA*, *ΔparZ*, *ΔparAZ* (**Fig. 4B, G, 6G-I**), consistent with the idea that ParA and ParZ works in the same pathway [7]. The absence of *parA* and/or *parZ* caused two major changes in chromosome folding: loci separated by 100 to 300 kb had increased interactions (**Fig. 6K-M**, red pixels below the black line) and loci separated by 300 kb or more had decreased interactions (**Fig. 6K-M**, blue pixels above the black line). Thus, ParA/ParZ acts to reduce mid-range (100-300 kb) and enhance long-range (>300 kb) DNA interactions on the chromosome. Since ParA/ParZ promotes chromosome segregation and spacing, we speculate that loss of ParA acting on DNA caused these changes in DNA interactions.

Finally, Δ*parAZBS,* which lacked both *parBS* and *parAZ,* formed its own group (group 6) (**Fig. 4B, H, 6J, N**). This mutant essentially exhibited an additive effect of *ΔparBS* (**Fig. 6C, F**) and *ΔparAZ* (**Fig. 6I, M**): decreased interactions below 150 kb (like in *ΔparBS*), increased mid-range (100-300 kb) interactions (as seen in *ΔparAZ*), and a complete loss of *terCL-terCR* interactions (**Fig. 6J, N**, black arrows). These effects can be explained by the independent actions of ParB/*parS* and ParA/ParZ that we discussed above.

Overall, our Hi-C analyses of these mutants indicate that the perturbation of genome interactions is correlated to the previously observed cytological defects in chromosome positioning and segregation [7]. Interestingly, although DNA interactions within the chromosome were changed in cells missing *parBS* or *parAZ*, the interactions between replicons (plasmid-chromosome and plasmid-plasmid interactions) remained similar to the control (**Fig. S5-S7**). Only in Δ*parAZBS*, plasmid-chromosome interactions were reduced, and plasmid-plasmid interactions were more evened out, which could be due to the entanglement of different copies of chromosomes in the polyploid cells [7].

## Discussion

In this study, we characterized the organization of the highly segmented genome of *B. burgdorferi* and the contribution of the chromosome partitioning proteins and SMC homologs to this organization. *B. burgdorferi* contains a linear chromosome and expresses an SMC protein, which is recruited by ParB/*parS* to the chromosomal origin like in many other bacteria. Notably, the *B. burgdorferi* chromosome does not have inter-arm interactions observed in other SMC-carrying bacteria [34, 36, 38, 39, 41, 45 , 46]. Nonetheless, SMC and its analog MksB contribute to long-range DNA contacts possibly through DNA looping. Interestingly, the absence of ParB/*parS* enhances SMC’s loop forming ability, suggesting that SMCs that load non-specifically outside of the chromosomal origin regions are more productive at forming DNA loops, while SMCs recruited by ParB to the origin is less so. Since *B. burgdorferi* is lacking ScpB and MksEF to form complete SMC and Mks complexes, it is possible that the loop formation mechanism by the incomplete complexes is different from the loop-extrusion activity of the holocomplexes [51–55]. For instance, it is possible that SMC or MksB alone can only facilitate long-range loop formation by binding to and bridging two DNA segments that are already in proximity.

The *B. burgdorferi* strain used in this study contains 18 plasmids. These plasmids showed differential interactions with the chromosome. Namely, plasmids lp17, lp21, lp25, and lp28-3 formed specific interactions with the chromosome at the *oriC* region, but the other 14 plasmids did not (**Fig. 3A, S6**). This pattern was highly reproducible in different mutants (**Fig. S5, S6**), suggesting that these plasmid-chromosome interactions are real, specific interactions. What are the molecular mechanism and biological function of these interactions? In *A. tumefaciens,* the secondary replicons cluster with the primary replicon at their origin regions through interactions between ParB homologs [41, 42], which prevents the loss of the secondary replicons [42]. In *B. burgdorferi*, we note that these interactions did not require ParB/*parS* or ParA/ParZ (**Fig. S5, S6**), suggesting that the molecular mechanism for these interactions is different from the centromeric clustering observed in *A. tumefaciens*. Although it is still possible that the four plasmids that interact with the chromosome may “piggyback” the chromosome to facilitate their own segregation and maintenance, it is also possible that these plasmid-chromosome interactions have functions unrelated to plasmid segregation. Indeed, 14 out of 18 plasmids did not interact with the chromosome origin, indicating that *B. burgdorferi* plasmids segregate largely independently from the chromosome. Notably, *B. burgdorferi* is polyploid with unequal copy number for each replicon [7] while *A. tumefaciens* newborn cells are haploid [41]. We postulate that the difference in ploidy might be one underlying factor accounting for the difference in organizing strategies between these two species. Our findings suggest that different species might take diverse strategies to organize and maintain segmented genomes.

The interactions between the plasmids on average are more frequent than plasmid-chromosome interactions and long-range intra-chromosomal interactions (**Fig. 1B, 2**). Interestingly, we observed all seven cp32 plasmids interact more frequently with one another, and cp26 and the ten linear plasmids preferentially interact with one another (**Fig. 3B**). This grouping does not seem to be correlated with plasmid size or copy number (**Fig. 1A, 3B**), and the mechanism for these preferential interactions remains to be explored.

Unlike in other bacteria studied to date, in *B. burgdorferi*, there are two partitioning system pairs, ParA/ParZ and ParB/*parS*, which co-regulate the spacing of the *oriC* copies in the cell. ParA/ParZ plays a more important role than ParB/*parS*. While removing ParB/*parS* only caused very mild defects in maintaining *oriC* spacing in the presence of ParA/ParZ, deleting both *parA* and *parBS* further disrupted the spacing pattern [7]. By Hi-C, we observed a similar trend in genome reorganization in these mutants: removing *parAZ* caused a significant increase of the medium-range (100-300 kb) interactions but double deletion of *parAZ* and *parBS* led to an additive increase in these interactions. Thus, the segregation defect is correlated with increased mid-range genome interactions. The causal relationship between chromosome segregation and genome folding is unclear and remains to be examined. We speculate that the tension exerted through the partitioning system leads to the change in DNA folding over the length of the chromosome, which in our case is the decrease of DNA interactions in the 100-300 kb range.

Despite the absence of inter-arm interactions on the chromosome, the two ends of the linear chromosome *terCL* and *terCR* interact, which requires ParA/ParZ and ParB/*parS*. The contribution of ParA/ParZ and ParB/*parS* to *terCL*-*terCR* interactions might be through different mechanisms. ParA/ParZ is required for the spacing of *oriC* copies [7]. Thus, it is possible that mis-positioning of chromosome copies reduces the frequency of *terCL-terCR* contacts. For ParB/*parS*, although it does not contribute much to the spacing of chromosome copies [7], it recruits SMC to the origin. Since SMC reduced *terCL-terCR* contacts (**Fig. 5F**), it is possible that ParB-mediated recruitment of SMC to the *oriC*-proximal *parS* site and away from chromosome arms lifts SMC’s inhibitory role in *terCL-terCR* interactions.

Altogether, our study identifies intrachromosomal, chromosome-plasmid, and plasmid-plasmid interactions of the most segmented bacterial genome known to date. We explored the contribution of SMC-family proteins and two partitioning systems to the folding and interactions of the genome. Although the exact mechanism for replicon interactions remains to be investigated, our study presents one step forward in the understanding of multipartite genome architecture and maintenance.

## Materials and methods

### General Methods

The *B. burgdorferi* strains used in this study are listed in **Table S1**. Cells were grown in exponential growth in complete Barbour-Stoenner-Kelly (BSK)-II liquid medium at 34°C in a humidified incubator and under 5% CO2 atmosphere [56, 57]. Complete BSK-II medium contained 50 g/L bovine serum albumin (Millipore, Cat. 810036), 9.7 g/L CMRL-1066 (US Biological, Cat. C5900-01), 5 g/L Neopeptone (Difco, Cat. 211681), 2 g/L Yeastolate (Difco, Cat. 255772), 6 g/L HEPES (Millipore, Cat. 391338), 5 g/L glucose (Sigma-Aldrich, Cat. G7021), 2.2 g/L sodium bicarbonate (Sigma-Aldrich, Cat. S5761), 0.8 g/L sodium pyruvate (Sigma-Aldrich, Cat. P5280), 0.7 g/L sodium citrate (Fisher Scientific, Cat. BP327), 0.4 g/L N-acetylglucosamine (Sigma-Aldrich, Cat. A3286), 60 mL/L heat-inactivated rabbit serum (Gibco, Cat.16120), and had a pH of 7.60. When noted, the following antibiotics were used: gentamicin at 40 μg/mL, streptomycin at 100 μg/mL, and kanamycin at 200 μg/mL [58–60]. Lists of strains, plasmids, oligonucleotides and Next-Generation-Sequencing samples can be found in Tables S1-S4.

### Growing cells for Hi-C

For Hi-C biological replicates, pairs of 100 mL cultures of each strain were inoculated and grown for two or three days. The cultures were fixed by addition of 37 mL 37% formaldehyde (Sigma-Aldrich, Cat. F8775) followed by rocking at room temperature for 30 min. Formaldehyde was inactivated using 7 mL 2.5 M glycine and rocking for 5 min. The samples were chilled on ice for 10 min, then pelleted at 4°C and 4,300 x g for 30 min in an Allegra X-14R centrifuge (Beckman Coulter) equipped with a swinging bucket SX4750 rotor. The pellet was resuspended in 1 mL ice-cold HN buffer (50 mM NaCl, 10 mM HEPES, pH 8.0) [61], then pelleted at 4°C and 10,000 x g for 10 min. The pellet was resuspended in 400 μL cold HN buffer, and 100 μL aliquots were frozen in a dry ice ethanol bath then stored at below -80°C.

### Hi-C

The detailed Hi-C procedure for *B. burgdorferi* was adapted from previously described protocols in *B. subtilis* [34] and *A. tumefaciens* [41]. Briefly, 5×10^8^ *B. burgdorferi* cells were used for each Hi-C reaction. Cells were lysed using Ready-Lyse Lysozyme (Epicentre, R1802M) in TE for 60 min, followed by 0.5% SDS treatment for 30 min. Solubilized chromatin was digested with DpnII and incubated for 2 hours at 37°C. The digested chromatin ends were repaired with Klenow and Biotin-14-dATP, dGTP, dCTP, dTTP. The repaired products were ligated in dilute reactions by T4 DNA ligase at 16°C overnight (about 20 hrs). Ligation products were reverse-crosslinked at 65°C overnight (about 20 hrs) supplemented with EDTA, 0.5% SDS and proteinase K. The DNA was then extracted twice with phenol/chloroform/isoamylalcohol (25:24:1) (PCI), precipitated with ethanol, and resuspended in 40 µl 0.1XTE buffer. Biotin at non-ligated ends was removed using T4 polymerase (4 hrs at 20°C) followed by extraction with PCI. The DNA was then resuspended in 105 μl ddH_2_O and sheared by sonication for 12 min with 20% amplitude using a Qsonica Q800R2 water bath sonicator. The sheared DNA was used for library preparation with the NEBNext UltraII kit (E7645) following the manufacturer’s instructions for end repair, adapter ligation, and size selection. Biotinylated DNA fragments were purified using 5 µl streptavidin beads following the manufacturer’s instructions. All DNA-bound beads were used for PCR in a 50 µl reaction for 14 cycles. PCR products were purified using Ampure beads (Beckman, A63881) and sequenced at the Indiana University Center for Genomics and Bioinformatics using NextSeq 500. Paired-end sequencing reads were mapped to the genome file of *B. burgdorferi* B31 (NCBI Reference Sequence GCA_000008685.2 ASM868v2) using the default setting with MAPAQ30 filter of Distiller (https://github.com/open2c/distiller-nf). Plasmids are arranged in this order: cp26, cp32-1, cp32-3, cp32-4, cp32-6, cp32-7, cp32-8, cp32-9, lp17, lp21, lp25, lp28-1, lp28-2, lp28-3, lp28-4, lp36, lp38 and lp54. Plasmids cp9, lp5 and lp56 are absent from our strain. The *B. burgdorferi* B31 genome was divided into 5-kb bins. Subsequent analysis and visualization were done using R and Python scripts.

### Hi-C analysis

The mapped Hi-C contact frequencies were stored in multi-resolution cooler files [62] and the Hi-C matrices were balanced using the iterative correction and eigenvector decomposition method [47]. The iterative correction method is a standard way to balance the Hi-C map such that the rows and columns sum to a constant value (typically 1), which helps to correct for biases in genomic coverage (e.g. how easy it is to capture or amplify specific genome regions). The iterative correction process and intuition for the procedure can be approximately summarized as follows: each individual value within a row is divided by the sum of values for that row to achieve a sum of 1 for every row. However, this normalization of the rows breaks the required symmetry of the Hi-C matrix. Therefore, row normalization is followed by column normalization where each individual value in a column is divided by the resulting sum of values for that column, which subsequently “unbalances” the rows and the row sum is no longer 1. As such, the process can be iteratively repeated until the row and column sums converge to 1 within a pre-defined error tolerance. This results in a balanced Hi-C matrix in which genomic coverage biases are minimized. We described the process starting with normalization of rows followed by columns. However, the procedure could equally have been applied by starting with columns instead of rows since the Hi-C matrix is symmetric about the primary diagonal. Unless otherwise specified, all Hi-C plots and downstream analyses were performed with this iterative correction.

Plots were generated with R or Python 3.8.15 using Matplotlib 3.6.2 [63]. Data were retrieved for plotting at 5-kb resolution. Pc(s) curves show the averaged contact frequency between all pairs of loci on the chromosome separated by set distance (s). The x-axis indicates the genomic distance of separation in kb. The y-axis represents averaged contact frequency in a logarithmic scale. The curves were computed for data binned at 5 kb. For the log_2_ ratio plots, the Hi-C matrix of each mutant was divided by the matrix of the control. Then, log_2_(mutant/control) was calculated and plotted in a heatmap using R.

### Clustering of strains based on Hi-C data

Clustering of strains based on the contact probability curves was done using the scikit-learn 1.1.3 k-means algorithm [50]. To determine the optimal number of clusters, we maximized the average Silhouette score. The silhouette score, *s(i)* is a metric that determines, for some collection of objects {i}, how well each individual object, *i*, matches the clustering at hand [64]. In our case, the collection of objects were the log-transformed contact frequency Pc(s) curves, which were computed as the average value of the contact frequency of pairs of loci separated by a fixed genomic distance. Average silhouette scores were computed for data clustered using k-means with varying the number of clusters ranging from 2 to 21. We found that the number of clusters that maximized the average silhouette score was 6, suggesting that 6 is the optimal number of clusters in the data.

### Generating expected plasmid-plasmid interaction frequencies map

Expected plasmid-plasmid interaction frequencies were computed using either copy number of the plasmids alone, as obtained by marker frequency analysis, or in combination with information on the plasmid lengths (**Fig. 1A**).

For the simulated plasmid-plasmid contact map using both the copy numbers and plasmid lengths (**Fig. S1A**), we first multiplied the average plasmid copy number relative to the *oriC* (i.e. which have values ranging between 0.5 and 1.4, see **Fig. 1A**) by the plasmid lengths in numbers of 5-kb bins (i.e. which have values between 3 and 10 bins per plasmid, see **Fig. 1A**) and rounded the resulting number to the nearest integer, *n_p_* for each plasmid *p*. The values of *n_p_* ranged between 2 and 14, and the total sum over all the plasmids, *p*, was N =∑_*p*_ 𝑛_*p*_ = 80. The simulated plasmid-plasmid “contact frequency” matrix was computed using the probability of randomly drawing a given pair of plasmids. The probability for drawing a plasmid, *p*, is *n_p_*/N. The resulting probability matrix from this calculation can be seen in **Fig. S1A** (top panel). To best compare the simulated plasmid-plasmid contact probability map with the experimental Hi-C data, we applied the iterative correction procedure [47] to this map. The resulting matrix is shown both with the same scale bar as the experimental Hi-C map (**Fig. S1A**, middle panel) and with a very fine color scale (**Fig. S1A**, bottom panel). We note that the iterative correction scheme tends to minimize the effects of copy number variation from one genome segment to another and this is why the expected (i.e. simulated) plasmid-plasmid contact map looks largely uniform when plotted with the same dynamic range as experimental data (**Fig. 3B, S1)**.

The simulated plasmid-plasmid contact map computed using only copy numbers was made in a similar fashion (**Fig. S1B**). For this method, instead of multiplying copy number by the length of the plasmid, a fixed integer number was used (in our case, 10) to convert the relative ratios into integer numbers. The method of computation was the same as that described above.

We make two important assumptions for this calculation: 1) plasmids constitute independent units of interaction, and 2) these independent units are “well mixed”. The independence of contacts assumption implies there are no restrictions on how many DNA segments may be simultaneously in contact with one another within a “Hi-C contact volume” and the identity of the DNA segments in contact does not matter. The “well mixed” assumption stipulates that independent DNA segments interact with equal probability with other DNA segments. Together, these assumptions allow us to compute the plasmid-plasmid interaction frequencies while safely ignoring other types of contacts such as plasmid-chromosome and chromosome-chromosome contacts.

### Plasmid construction

Plasmid pΔmksB(gent) was generated in the following manner: (i) nucleotides 874996 through 876527 of the B31 chromosome were PCR-amplified with primers NT968 and NT969; (ii) the gentamicin cassette of pKIGent_parSP1_phoU [7] was PCR-amplified with primers NT970 and NT971; (iii) nucleotides 879168 through 880691 of the B31 chromosome were PCR-amplified with primers NT972 and NT973; (iv) the suicide vector backbone of pΔparA(kan) [7] was PCR-amplified with primers NT974 and NT975; and (v) the four PCR fragments listed above were digested with DpnI (New England Biolabs), gel-purified, and subjected to Gibson assembly [65] using New England Biolabs’ platform. The assembled plasmid was introduced into *Escherichia coli* strain NEB 5-alpha (New England Biolabs) by heat shocking. The resulting strain (CJW7512) was grown at 30°C on LB plates or in Super Broth liquid medium with shaking, while 15 μg/mL gentamicin was used for selection.

### Strain construction

To generate strain CJW_Bb605, 75 μg of plasmid pΔmksB(gent) were digested with ApaLI (New England Biolabs) in a 500 μL reaction volume for 4 hours. The DNA was then ethanol precipitated [66], dried, and resuspended into 25 μL sterile water. The resulting DNA suspension was then electroporated at 2.5 kV, 25 μF, 200 Ω, 2 mm-gap cuvette [67, 68] into 100 μL of electrocompetent cells made [69] using *B. burgdorferi* strain S9. The electroporated bacteria were transferred immediately to 6 mL BSK-II medium and allowed to recover overnight at 34°C. The next day, a fraction of the culture was embedded in 25 mL of semisolid BSK-agarose medium containing gentamicin per 10-cm round Petri dish, as previously described [70]. The semisolid BSK-agarose mix was made by mixing 2 volumes of 1.7% agarose in water, sterilized by autoclaving, then melted and pre-equilibrated at 55°C, with 3 volumes of BSK-1.5 medium, which was also equilibrated at 55°C for at most 5 minutes. BSK-1.5 contained 69.4 g/L bovine serum albumin, 12.7 g/L CMRL-1066, 6.9 g/L Neopeptone, 3.5 g/L Yeastolate, 8.3 g/L HEPES, 6.9 g/L glucose, 6.4 g/L sodium bicarbonate, 1.1 g/L sodium pyruvate, 1.0 g/L sodium citrate, 0.6 g/L N-acetylglucosamine, and 40 mL/L heat-inactivated rabbit serum, and had a pH of 7.50. After 10 days of growth in the BSK-agarose semisolid matrix, an individual colony was expanded in liquid culture and confirmed by PCR to have undergone correct double crossover homologous recombination of the suicide vector, thus yielding strain CJW_Bb605. This strain was also confirmed by multiplex PCR [71] to contain all endogenous plasmids contained by its parent.

Further information and requests for strains, plasmids, resources, reagents, and analytical scripts should be directed to and will be fulfilled by the corresponding authors with appropriate Material Transfer Agreements.

## Acknowledgements

We thank the Wang and Jacobs-Wagner labs for discussions and support, the Indiana University Center for Genomics and Bioinformatics for assistance with high-throughput sequencing. Support for this work comes in part from the Pew Innovation Fund (C.J.-W.), and the National Institutes of Health R01GM141242 and R01GM143182 (X.W.). This research is a contribution of the GEMS Biology Integration Institute, funded by the National Science Foundation DBI Biology Integration Institutes Program, Award #2022049 (X.W.). Christine Jacobs-Wagner is an investigator of the Howard Hughes Medical Institute.

## Supplemental Information

Supplemental information includes seven figures and four tables.

## Author Contributions

Z.R., C.N.T., C.J.-W. and X.W. designed the study. Z.R. and X.W. performed Hi-C experiments and analyses. C.N.T. generated plasmids and strains and collected cells for Hi-C experiments. H.B.B. developed methods for analysis and generated figure plots. C.J.-W. and X.W. supervised the project and acquired funding. Z.R. and X.W. wrote the manuscript with input from all authors.

## Supplementary Information for

**This PDF file includes:**

Tables S1 to S4

SI References

---

**Figure S1.**
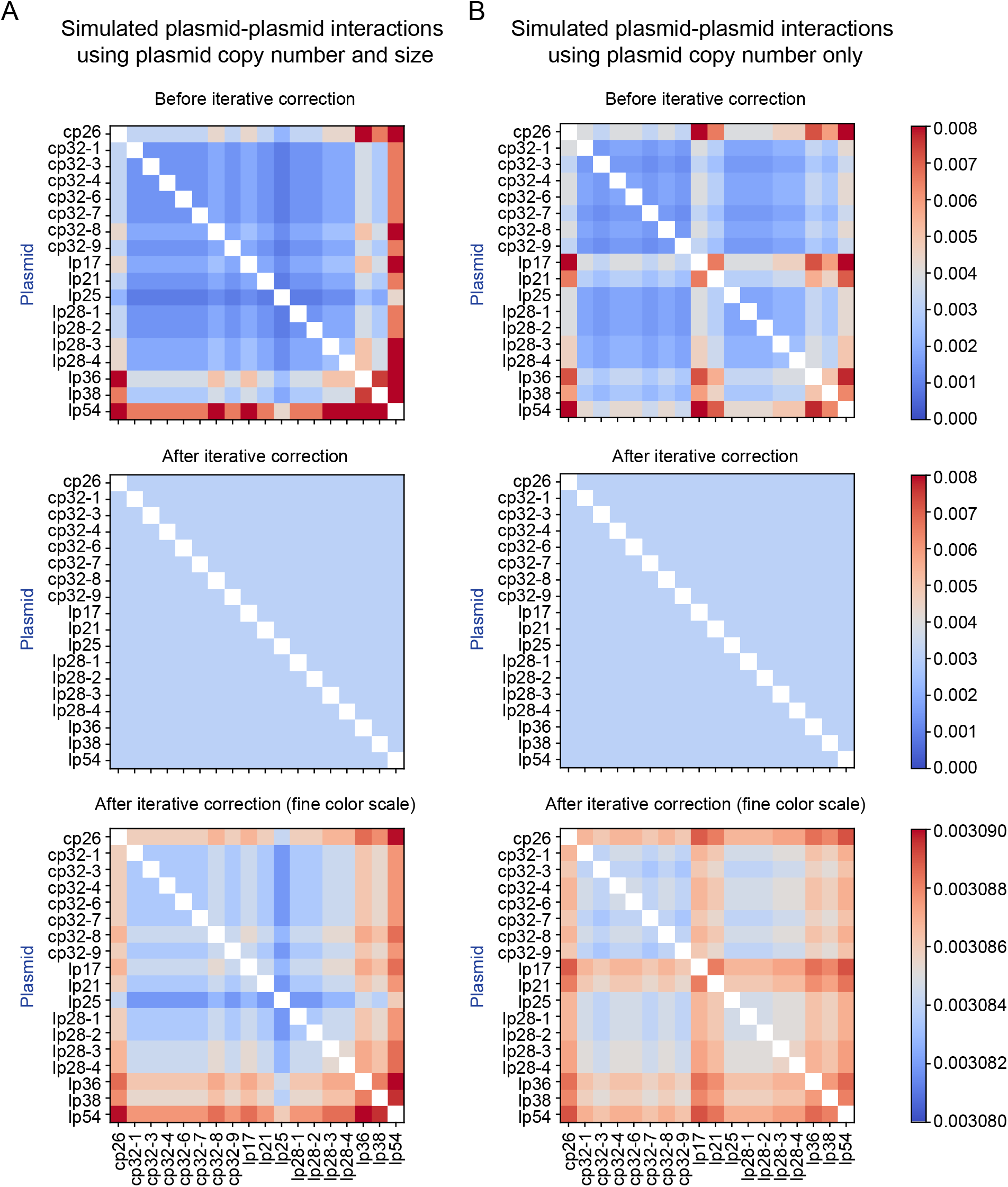
Simulated plasmid-plasmid interaction frequency. The expected contact probability between plasmids was calculated under the assumptions that plasmids are independent of one another and are “well mixed” within the cytoplasm. The calculation was performed using copy number and plasmid length together (**A**) or using only plasmid copy numbers (**B**). Top panels, the exact contact frequency expected between plasmid segments. Middle panels, the contact frequency expected between plasmids after application of the iterative correction normalization procedure. Bottom panels, the same as middle panels, but shown with a much finer color scale. The color scale depicting contact frequency in arbitrary unit is shown at the right. We note that the residual resemblance between bottom and top panels results from the fact that the iterative correction procedure only asymptotically approaches 1 (see Materials and Methods).

**Figure S2.**
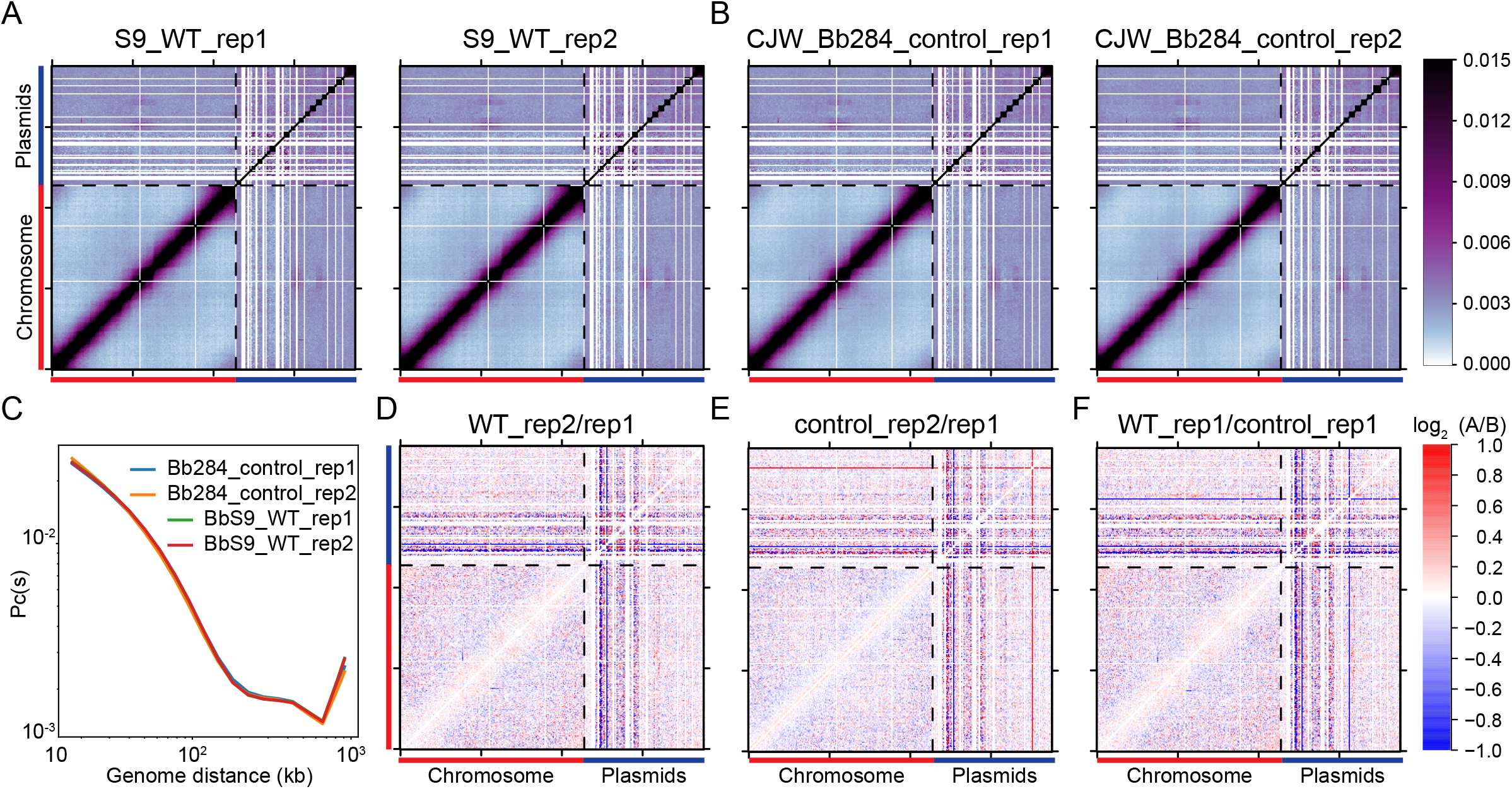
Comparison of WT and control. **(A-B)** Normalized Hi-C interaction maps of *B. burgdorferi* strains S9 (WT) and the control strain CJW_Bb284. Two biological replicates of each strain (rep1 and rep2) are shown. The color scale depicting Hi-C interaction scores in arbitrary unit is shown at the right. We note that *PflaB-aadA* sequence from the chromosome is inserted in *bbe02* region lp25. Short-range intra-chromosomal interactions involving the *flaB* promoter region could be assigned to lp25 and account for the interactions between lp25 and the promoter region of *flab* on the chromosome at ∼150 kb. **(C)** Pc(s) curves of the four samples. Pc(s) curves show the averaged contact frequency between all pairs of loci on the chromosome separated by set distance (*s*). The x-axis indicates the genomic distance of separation in kb. The y-axis represents averaged contact frequency. The curves were computed for data binned at 5 kb. **(D-F)** Log_2_ ratio plots comparing different Hi-C matrices. Log_2_(matrix 1/matrix 2) was calculated and plotted in the heatmaps. Matrix 1 / matrix 2 are shown at the top of each plot. The color scale is shown at the right of panel **(F)**.

**Figure S3.**
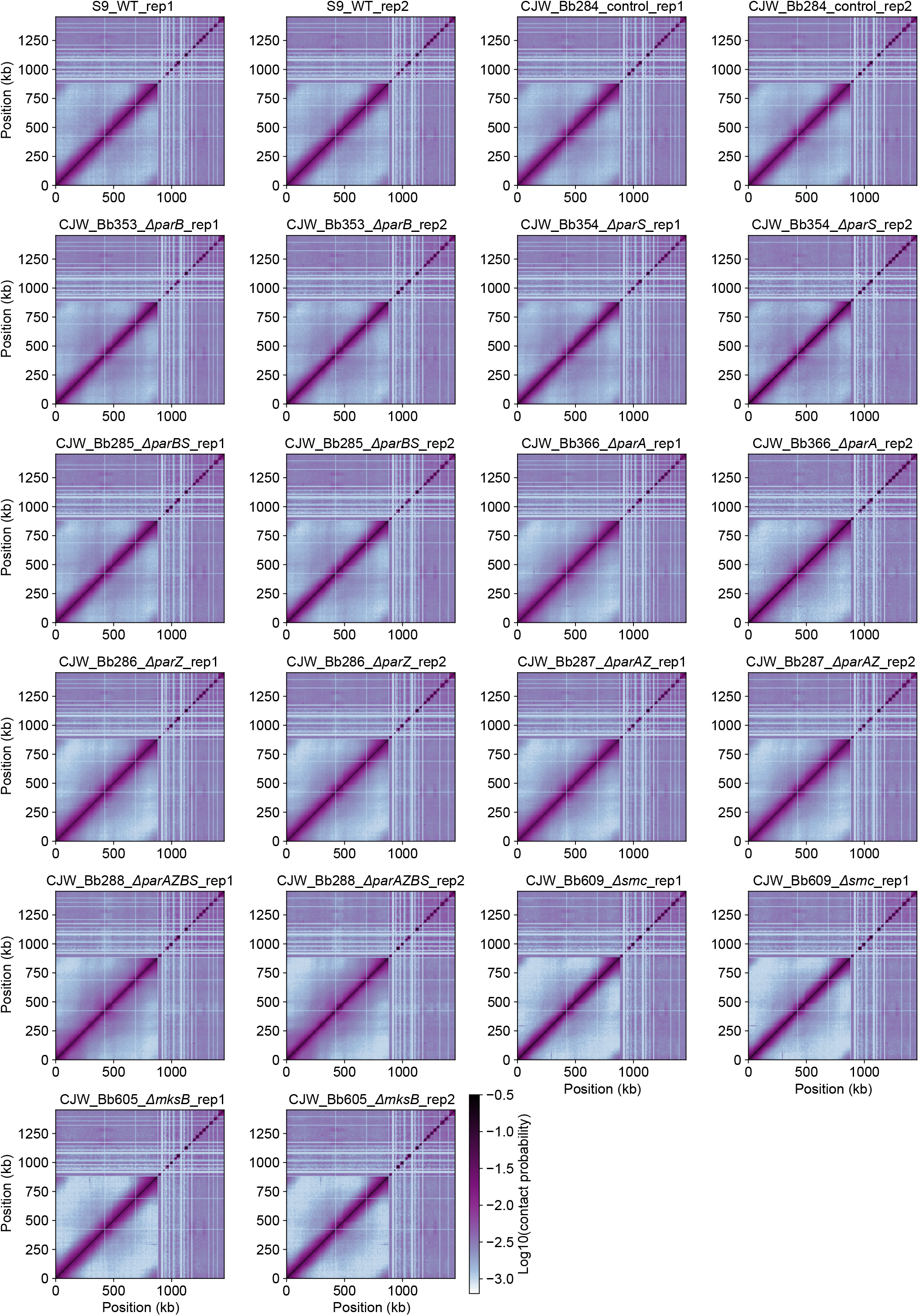
Hi-C samples used in this study. The normalized Hi-C plots of all the 22 experiments. The color scale depicting Hi-C interaction scores is shown in log_10_.

**Figure S4.**
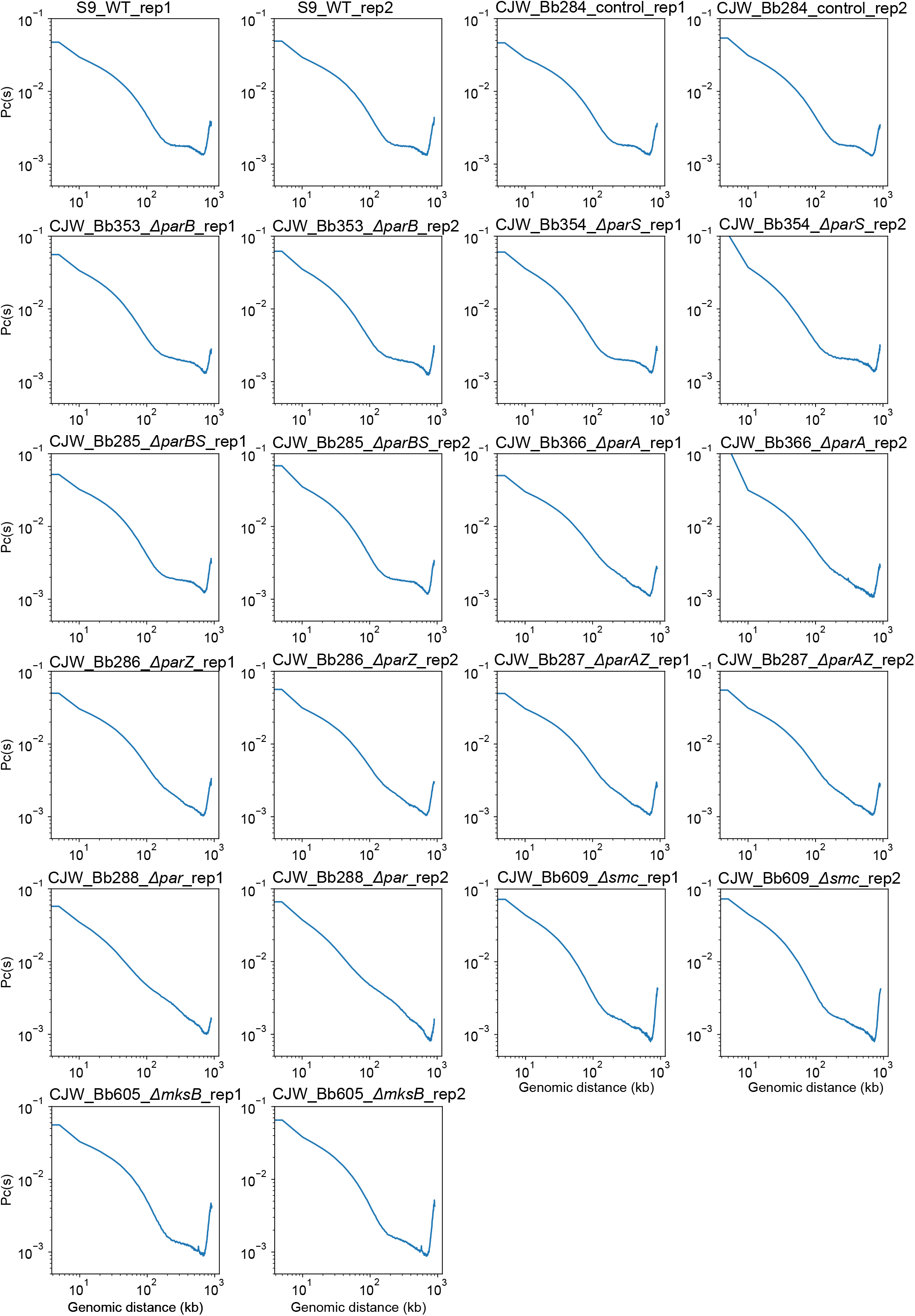
Individual Pc(s) curves of all the samples analyzed in this study. Pc(s) curves of all the 22 Hi-C experiments. x-axis indicates genomic distance and y-axis shows averaged contact frequency.

**Figure S5.**
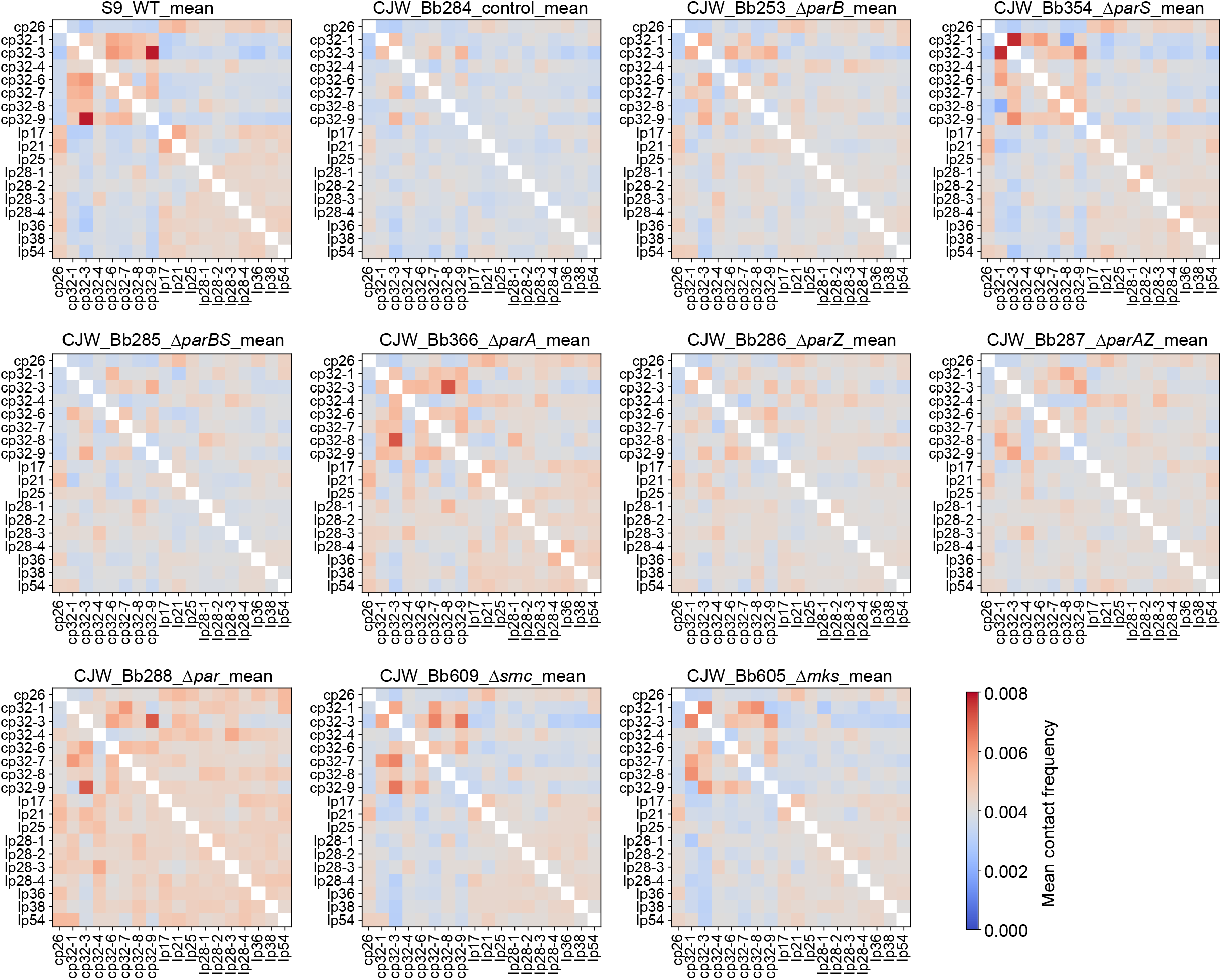
Plasmid-chromosome interactions in different mutants. Heatmap of plasmid-chromosome interaction frequencies are shown. The x-axis shows chromosome location in kb. The y-axis specifies the different plasmids analyzed. The color indicates the contact frequency between plasmid and chromosome loci. Each graph plots the mean value of two biological replicates found in **Fig. S3**. Data are binned at 5-kb resolution.

**Figure S6.**
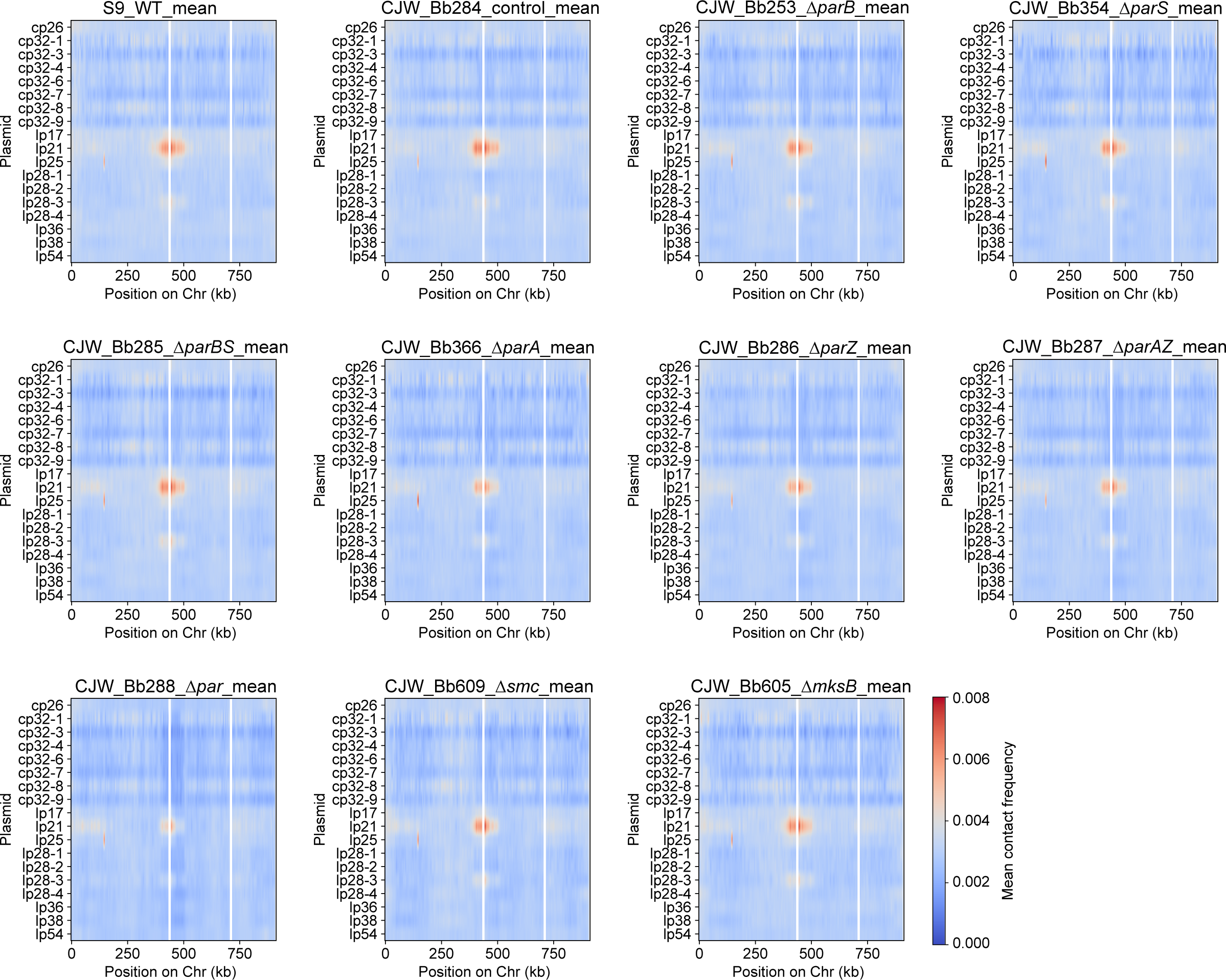
Plasmid-chromosome interactions in different mutants organized by plasmids. Heatmaps of plasmid-chromosome interaction frequencies are shown. The x-axis shows the chromosome location in kb. The y-axis specifies the different mutants. The color indicates the contact frequency between plasmid and chromosome loci. Each graph plots the mean value of two biological replicates found in **Fig. S3**. Data are binned at 5-kb resolution.

**Figure S7.**
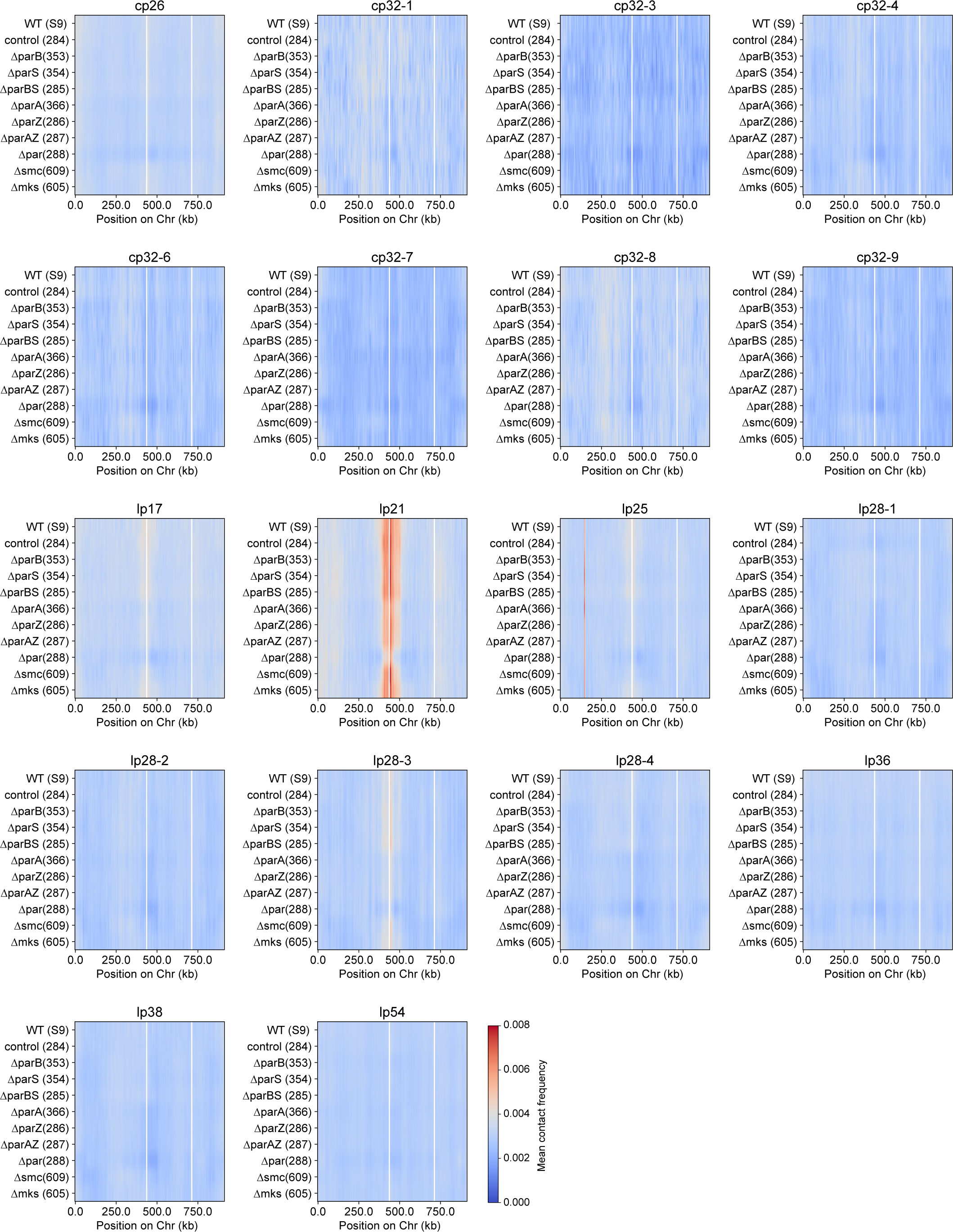
Plasmid-plasmid interactions in different mutants. Plasmid-plasmid contact frequencies in different strains. The x and y axes indicate the plasmids analyzed. The color shows the computed contact frequency. Each graph plots the mean of two biological replicates found in **Fig. S3**. Data are normalized such that the sum of each row has the same total score.

**Table S1.** Bacterial strains used in this study.

| Strain | Genotype | Antibiotic resistance | Reference | Figure |
| --- | --- | --- | --- | --- |
| S9 | Transformable derivative of the *B. burgdorferi* type strain B31; lacks endogenous plasmids cp9, lp5, and lp56; also known as B31-A3-68-$\Delta bbe02::PflaB-aadA$ | Sr | [1] | 1-4, S2-S7 |
| CJW_Bb284 | S9-derived control strain; has gentamicin resistance cassette inserted between *parZ* and *parB* | Sr, Gm | [2] | 4A-C, 5A, 5D-I, S2-S7 |
| CJW_Bb285 | S9-derived $\Delta parBS$ strain | Sr, Gm | [2] | 4ABF, 6CF, S3-7 |
| CJW_Bb286 | S9-derived $\Delta parZ$ strain | Sr, Gm | [2] | 4ABG, 6HL, S3-7 |
| CJW_Bb287 | S9-derived $\Delta parAZ$ strain | Sr, Gm | [2] | 4ABG, 6IM, S3-7 |
| CJW_Bb288 | S9-derived $\Delta parAZBS$ strain | Sr, Gm | [2] | 4ABH, 6JN, S3-7 |
| CJW_Bb353 | S9-derived $\Delta parB$ strain | Sr, Gm | [2] | 4ABF, 6AD, S3-7 |
| CJW_Bb354 | S9-derived $\Delta parS$ strain | Sr, Gm | [2] | 4ABF, 6BE, S3-7 |
| CJW_Bb366 | S9-derived $\Delta parA$ strain | Sr, Km | [2] | 4ABG, 6GK, S3-7 |
| CJW_Bb605 | S9-derived $\Delta mksB$ strain | Sr, Gm | This study | 4ABE, 5BEH, S3-7 |
| CJW_Bb609 | S9-derived $\Delta smc$ strain | Sr, Gm | [2] | 4ABD, 5CFI, S3-7 |

Sr, streptomycin resistance; Gm, gentamicin resistance; Km, kanamycin resistance.

**Table S2.**
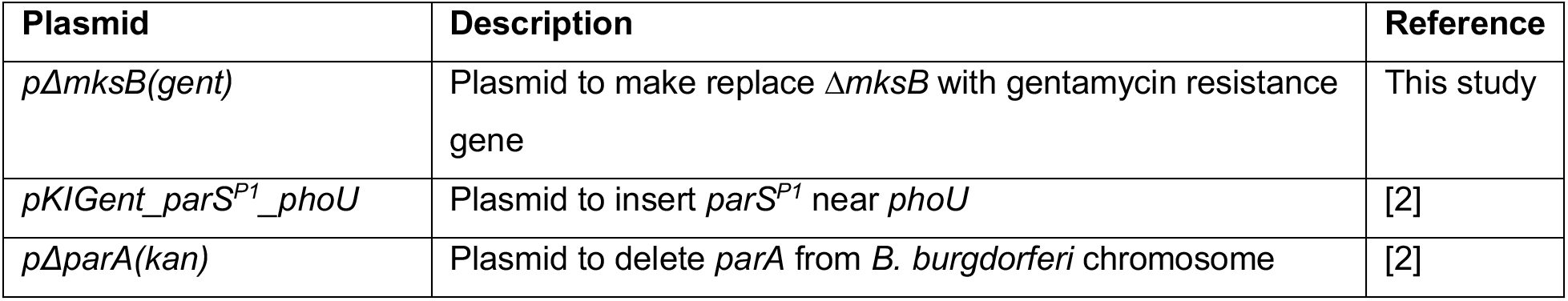
Plasmids used in this study.

**Table S3.**
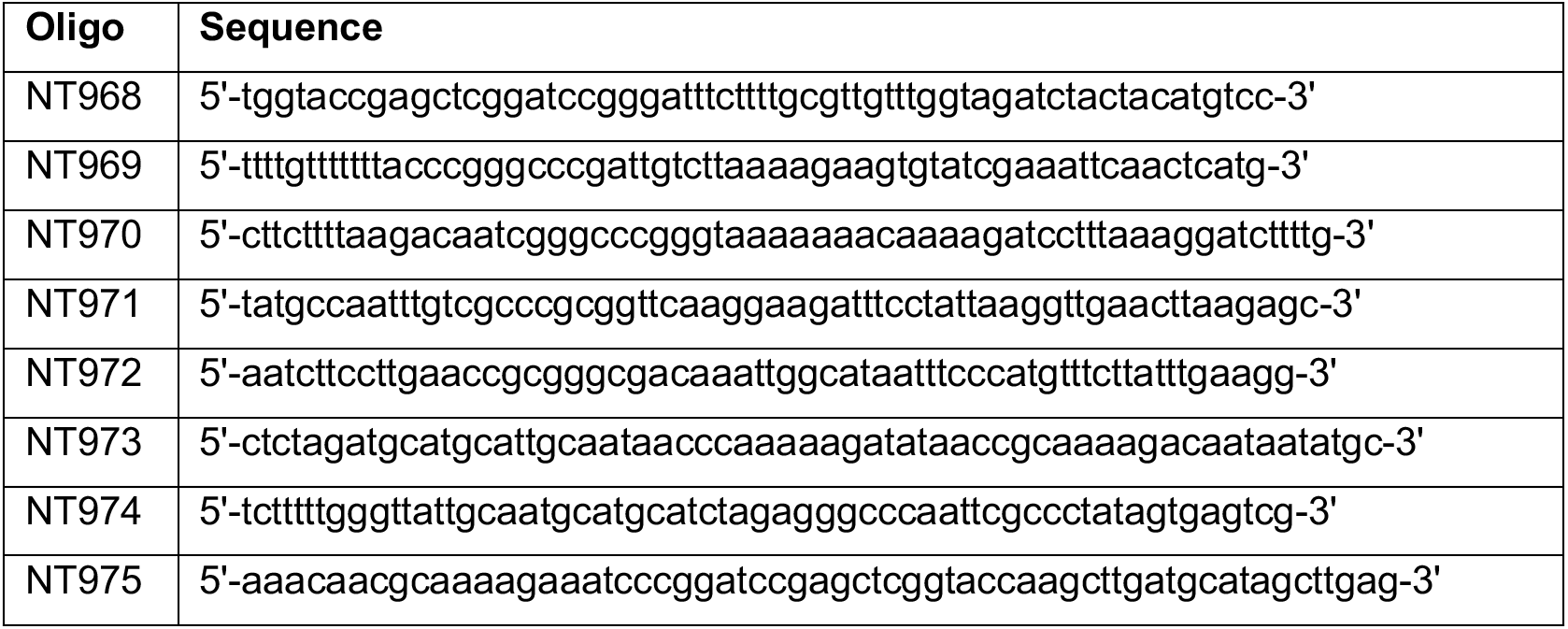
Oligonucleotides used in this study.

**Table S4.** Next generation sequencing samples used in this study.

| **Sample name** | **Figure** | **Reference** |
| --- | --- | --- |
| HiC_CJW_Bb284_rep1 | 4A-C, 5A, 5D-I, S2-7 | This study |
| HiC_CJW_Bb284_rep2 | 4A-C, S2-7 | This study |
| HiC_CJW_Bb285_rep1 | 4ABF, 6CF, S3-7 | This study |
| HiC_CJW_Bb285_rep2 | 4ABF, S3-7 | This study |
| HiC_CJW_Bb286_rep1 | 4ABG, 6HL, S3-7 | This study |
| HiC_CJW_Bb286_rep2 | 4ABG, S3-7 | This study |
| HiC_CJW_Bb287_rep1 | 4ABG, 6IM, S3-7 | This study |
| HiC_CJW_Bb287_rep2 | 4ABG, S3-7 | This study |
| HiC_CJW_Bb288_rep1 | 4ABH, 6JN, S3-7 | This study |
| HiC_CJW_Bb288_rep2 | 4ABH, S3-7 | This study |
| HiC_CJW_Bb353_rep1 | 4ABF, 6AD, S3-7 | This study |
| HiC_CJW_Bb353_rep2 | 4ABF, S3-7 | This study |
| HiC_CJW_Bb354_rep1 | 4ABF, 6AD, S3-7 | This study |
| HiC_CJW_Bb354_rep2 | 4ABF, S3-7 | This study |
| HiC_CJW_Bb366_rep1 | 4ABG, 6GK, S3-7 | This study |
| HiC_CJW_Bb366_rep2 | 4ABG, S3-7 | This study |
| HiC_CJW_Bb605_rep1 | 4ABE, S3-7 | This study |
| HiC_CJW_Bb605_rep2 | 4ABE, 5BEH, S3-7 | This study |
| HiC_CJW_Bb609_rep1 | 4ABD, 5CFI, S3-7 | This study |
| HiC_CJW_Bb609_rep2 | 4ABD, S3-7 | This study |
| HiC_CJW_S9WT_rep1 | 1-4, S2-7 | This study |
| HiC_CJW_S9WT_rep2 | 4A-C, S2-7 | This study |

